# Characterization of Tumor-Associated Endothelial Cells and the Development of a Prognostic Model in Pancreatic Ductal Adenocarcinoma

**DOI:** 10.1101/2023.07.22.550139

**Authors:** Jun Wu, Yang Liu, Qi Fu, Zhi Cao, Xun Li, Xiaodong Ma

## Abstract

**Background:** Pancreatic ductal adenocarcinoma (PDAC) is a highly aggressive malignancy characterized by a complex tumor microenvironment. Angiogenesis is of paramount importance in the proliferation and metastasis of PDAC. However, currently, there are no well-defined biomarkers available to guide the prognosis and treatment of PDAC.

**Results:** In this study, we investigated the interactions between tumor-associated endothelial cells (TAECs) and tumor cells in PDAC, and identified a specific subset of TAECs characterized by high expression of COL4A1. COL4A1+ endothelial cells interact with tumor cells through the COLLAGEN signaling pathway to promote tumor cell proliferation, migration, and invasion. We also observed activation of HOXD9 in COL4A1+ endothelial cells. Based on these findings, we developed a prognostic model called TaEMS, which accurately predicts patient prognosis. TaEMS identified high-risk patients enriched in cell cycle-related pathways and low-risk patients enriched in focal adhesions, smooth muscle regulation, and immune pathways. Moreover, high-risk patients displayed a reduced level of immune cell infiltration, indicating the presence of a “cold tumor” phenotype.

**Conclusions:** Overall, our study uncovered an intricate crosstalk between TAECs and tumor cells in PDAC, emphasizing the role of HOXD9 and highlighting the potential of TaEMS as a prognostic biomarker for precise therapies.

## 1. Introduction

Pancreatic ductal adenocarcinoma (PDAC) is one of the most aggressive and lethal types of cancer and is known for its late detection, rapid disease progression, and limited therapeutic interventions[1, 2]. Despite significant advancements in cancer research, the prognosis of patients with PDAC remains challenging, emphasizing the need for a deeper understanding of the underlying mechanisms driving tumor development and progression[3].

Tumor angiogenesis, the formation of new blood vessels to support tumor growth, is a critical process in cancer biology[4]. It plays a crucial role in the supply of oxygen and nutrients to the tumor and facilitates metastasis in the tumor microenvironment[5]. Dysregulation of angiogenesis has been implicated in PDAC, contributing to tumor aggressiveness and treatment resistance[6–8]. Tumor vasculature is a complex system, and angiogenesis can provide escape routes and nutritional support for PDAC cancer cells[9]. Most patients had vascular infiltration and metastasis, and Chen et al. reported that PDAC cells can promote tumor angiogenesis under hypoxic conditions[10]. However, insights into tumor-associated endothelial cells in the PDAC tumor microenvironment are still lacking.

In recent years, single-cell sequencing technologies have revolutionized our ability to unravel cellular heterogeneity within tumors[11–13]. By dissecting the complex tumor microenvironment at the single-cell level, researchers have identified distinct cell populations and their molecular characteristics, shedding light on the intricate interplay between cancer, stromal, and immune cells[14–16].

In our study, we used single-cell data analysis to decipher the cellular landscape of PDAC and focused on a specific subtype of TAECs. Endothelial cells play a crucial role in angiogenesis and are closely associated with tumor growth and metastasis. Furthermore, we constructed a prognostic risk model based on the signatures of these TAECs. By integrating clinical data, we evaluated and validated the performance of this model. This research enhances our understanding of the tumor vasculature in PDAC, and the development of a prognostic risk model provides valuable patient stratification and treatment decision-making information for clinical practitioners.

## 2. Materials and methods

### 2.1 Data collection

Single-cell RNA-sequencing data of PDAC samples (GSE212966 and CRA001160) were obtained from the Gene Expression Omnibus (GEO, https://www.ncbi.nlm.nih.gov/geo/)[17] and Genome Sequence Archive (GSA, https://ngdc.cncb.ac.cn/gsa/) databases[18]. The bulk RNA-sequencing data of PDAC samples were obtained from TCGA, ICGC (https://dcc.icgc.org/) and GEO (GSE183795) databases.

### 2.2 Single-cell RNA-sequencing(scRNA-seq) data process and integration

To generate the feature-barcode gene expression matrix, the raw data (GSE212966) were aligned to the human reference genome (GRCh38 transcriptome, http://cf.10Xgenomics.com/supp/cell-exp/refdata-cellranger-GR <u>Ch38-1.2.0.tar.gz</u>) using Cell Ranger (version 3.0.1, 10X Genomics). We directly downloaded the gene expression matrix of CRA001160 provided by the GSA database. The scRNA-seq data, including GSE212966 and CRA001160, were analyzed using the R package Seurat v4.3.0 in R 4.1.1[19]. In GSE212966, the first step was to filter out low-quality cells with a cutoff value of less than 500 total feature RNA and more than 30 mitochondrial RNA. After normalizing gene expression in each cell, the data were integrated using SCTransform integration workflow (https://satijalab.org/seurat/articles/integrati <u>on_introduction.html</u>). A total of 2000 integration anchors representing the nearest neighbors within the datasets were identified.

### 2.3 Dimension reduction and cell clustering

To reduce dimensions, principal component analysis (PCA) was performed (npcs=50), followed by t-distributed stochastic neighbor embedding (tSNE). Then 38 clusters were found after applying the FindNeighbors (dims=1:30) and FindClusters (resolution=0.8) functions. Finally, we annotated a total of 47,671 cells across nine samples with 13 major cell clusters based on the average gene expression of well-known markers, including CD8+ T cells (*CD3D, CD8A*, and *CD8B*), fibroblasts (*COL1A1* and *DCN*), CD4+ T cells (*CD3D, CD4,* and *IL7R*), B cells (*CD79A* and *MS4A1*), macrophages(*CD68* and *MARCO*), endothelial cells (*PECAM1* and *VWF*), pericytes (*RGS5*, *ACTA2*, and *CSPG4*), dendritic cells (*PTPRC* and *CD1C*), epithelial cells (*EPCAM* and *KRT8*), monocytes (*CD14* and *S100A8*), NK cells (*NKG7* and *GNLY*), plasma cells (*CD79A*, *IGHA1*, and *IGLC2*), and mast cells (*CPA3* and *KIT*). In CRA001160, cell type references for each barcode were performed using GSA[14]. We annotated 57530 cells across 35 donors with 11 cell clusters in CRA001160 for validation.

### 2.4 Differential gene expression analysis

We further performed differential gene expression analysis using the “FindAllMarkers” function in Seurat to identify the differentially expressed genes (DEGs). The following parameters were set: logfc.threshold=0.25, only.pos=T, test.use=wilcox. The remaining parameters were set to their default values. The R package pheatmap was used to visualize the gene expression heatmap after scaling.

### 2.5 Transcription factor regulon analysis

The activity of regulatory regulons was determined using the R package SCENIC v1.3.1[20, 21]. Regulatory regulons were calculated using the AUCell module with default parameter settings. The background dataset was downloaded using the R package RcisTarget v1.20.0. Endothelial cell clusters and expression matrices obtained from Seurat were passed through the SCENIC pipeline, and the resulting transcriptional regulatory activity was visualized using heatmaps. The gene expression values corresponding to the transcriptional regulatory regulons were visualized using FeaturePlot and DotPlot functions in Seurat.

### 2.6 Prediction of endothelial cell differentiation and development

CytoTRACE is a powerful computational tool that predicts cell differentiation states based on scRNA-seq data[22]. Following the application of the CytoTRACE algorithm, each cell was assigned a stemness score to assess its degree of differentiation. To calculate the stemness scores for endothelial cells, we used the R package CytoTRACE v0.3.3. The CytoTRACE scores range from 0 to 1, where lower scores indicate higher differentiation and higher scores indicate greater stemness. Monocle2 is a computational framework used to analyze scRNA-seq data to reconstruct developmental trajectories and identify gene expression changes along these trajectories[23]. We then employed the R package Monocle2 v2.28.0 for pseudotime analysis of the 11 endothelial cell clusters, allowing us to project cell subtypes and stemness scores onto a pseudotime trajectory. This trajectory was constructed using the top 2000 differentially expressed genes, selected based on adjusted p-values using the FindAllMarkers function in Seurat software. Pseudotime was assigned based on the descending order of stemness scores obtained from CytoTRACE, providing insights into the temporal progression of cell differentiation.

### 2.7 Cell-cell communication analysis

To identify the communication between different cells, the R package CellChat v1.4.0 was used[24], which utilized paired receptor-ligand information stored in its internal dataset “CellChatDB.human”. We analyzed the signals between COL4A1+ endothelial cells and other cells using the official workflow (https://github.com/sqjin/CellChat). These included functions such as “identifyOverExpressedGenes”, “projectData”, and “projectData” with the parameter “min.cells=3”, “computeCommunProbPathway” and “aggregateNet”. Visualization of the ligand-receptor pairs was performed using the functions “netVisual_bubble”, “netAnalysis_computeCentrality”, “netVisual_heatmap”, and “plotGeneExpression”.

### 2.8 Construction and validation of prognostic signature

Identification of a set of highly expressed genes in tumor-associated COL4A1+ endothelial cells based on scRNA-seq. We used univariate Cox regression analysis to evaluate the prognostic significance of these genes in terms of overall survival among patients with PDAC in the TCGA-PAAD dataset. Genes with a significance level of p < 0.05 were identified as prognostic genes. We further refined the prognostic gene set using the R package glmnet v4.1-7, employing the least absolute shrinkage and selection operator (LASSO) Cox proportional hazards regression analysis method[25]. Using the “cv.glmnet” function, we performed ten-fold cross-validation to select the optimal model, resulting in a list of genes with non-zero coefficients (Supplementary Tables S2). Finally, we constructed a prognostic risk model by combining the gene mRNA expression levels with their corresponding risk coefficients. Based on the median, patients were divided into high-risk and low-risk groups. Patient information is shown in Supplementary Tables S1. The TaEMS calculation formula was developed as below:

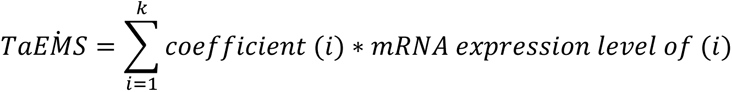

To validate the prognostic ability of the tumor-associated endothelial marker signature (TaEMS), we used the R package survminer v0.4.9 to determine statistical significance using the log-rank test and visualized the results using Kaplan-Meier plots. The R package timeROC v0.4 was used to calculate the area under the curve (AUC). Lastly, we validated the predictive ability of the model through survival analysis and AUC calculation using independent datasets from ICGC and GEO.

### 2.9 Cancer hallmark enrichment and GSEA analyses

We downloaded 50 tumor hallmark gene sets for Homo sapiens from the Molecular Signatures Database (MSigDB, https://www.gseamsigdb.org/gsea/msigdb). We utilized the function “gsva” from the R package GSVA v1.48.1[26] with the method set to “ssgsea” to score hallmark gene sets, and the scaled gene set scores were visualized using a heatmap.

For KEGG pathway enrichment analysis in the prognostic risk model, we obtained the Homo sapiens KEGG pathway gene sets from MSigDB. Gene set enrichment analysis (GSEA) was conducted using the R package fgsea v1.26.0, with the following parameters: minSize=5, maxSize=500, and nperm=1000. The genes were ranked according to the differential fold-change between the high-risk and low-risk groups.

### 2.10 Immune infiltration analysis

We used the R package ESTIMATE v1.0.13 to evaluate the levels of stromal and immune cell infiltration. Based on the mRNA sequencing data of PDAC samples from TCGA-PAAD, we calculated stromal and immune scores, ESTIMATE scores, and tumor purity scores for each sample. The samples were then divided into high and low-risk groups for comparison, and statistical significance was determined using the Wilcoxon test.

### 2.11 Analysis of antiangiogenesis drug treatment efficacy assessment

To assess whether there are differences in the predicted treatment response to different drugs between high-risk and low-risk groups, we used the R package pRRophetic v0.5 to predict mRNA expression matrices of PDAC samples from the TCGA-PAAD dataset. pRRophetic combines machine learning and statistical models to model and train on large-scale data sets to predict patient responses to various drugs. The function pRRopheticPredict was used with the parameters set as follows: tissueType=“all”, dataset=“cgp2014”. A total of 138 drugs were evaluated, and the results were visualized using the ggboxplot function in the R package ggpubr.

### 2.12 Predictive Immunotherapy in IMvigor210

To validate the predictive capability of TaEMS in immunotherapy, we used the IMvigor210 dataset. IMvigor210 is a research project that focuses on immunotherapy for bladder cancer[27]. It involves gene expression profiling of patient tumor tissues and statistical analysis of survival and treatment response. TaEMS is a prognostic model specifically developed for tumor-associated endothelial cells in PDAC, and it has also exhibited robust predictive capability in the IMvigor210 cohort.

### 2.13 Survival analysis

The prognostic significance of HOXD9 in patients with PDAC was assessed using the R package survival 3.5-5. The optimal threshold for grouping the expression values of HOXD9 was determined using the function “maxstat.test” from the R package “maxstat” with the parameter “smethod” set to “LogRank”. Subsequently, the results were visualized using the “ggsurvplot” function in the R package survminer v0.4.9.

### 2.14 Immunofluorescence staining

PDAC tissue samples preserved in formalin were subjected to standard dehydration and paraffin embedding procedures. The paraffin blocks were then cut into 5 µm-thick slides and mounted onto glass slides. Subsequently, the paraffin-embedded sections underwent dewaxing, rehydration, blockade of endogenous peroxidase activity, and high-temperature antigen retrieval. The sections were subsequently prepared for multiplex immunofluorescence (IF) staining assays. Sections were incubated overnight at 4°C with the following primary antibodies: anti-HOXD9 (Santa Cruz Biotechnology, Inc.; cat. no. sc-137134; 1:100), anti-CD31 (rabbit antibody; ABclonal; cat. no. A19014; 1:300) and anti-EPCAM (rabbit antibody; cat. no. A19301; 1:300). 4’,6-Diamidino-2-phenylindole (DAPI) was used to stain the cell nucleus. After washing, the corresponding Alexa Fluor secondary antibodies were applied for 1 hour in the dark. The stained samples were imaged using a Zeiss LSM 900 laser scanning confocal microscope with excitation/emission filters for Alexa Fluor 488, 568, and 647, and DAPI. Optical sections were obtained for each channel. Image analysis was used to quantify the tissue localization and expression levels of EpCAM, CD31, and nuclear HOXD9. DAPI staining allowed visualization of the total nuclei. This multiplexed immunofluorescence staining combined with confocal imaging enabled simultaneous detection of epithelial cells, vasculature, HOXD9 expression, and total nuclei within intact tissue sections.

### 2.15 Statistical analysis

When appropriate, statistical differences between groups were evaluated using Student’s t test or Wilcoxon test, depending on the data distribution. Survival analysis was performed using the log-rank test. A p value less than 0.05 was considered statistically significant and denoted as *, while p < 0.01, represented as **, p < 0.001 as ***, and p < 0.0001 or below as ****.

## 3. Results

### 3.1 Workflow and Landscape of Cell Composition in PDAC

To identify tumor-associated endothelial cells and their functions in PDAC, we collected multiple datasets from TCGA, GEO, GSA, and ICGC databases for this study and performed integrated analysis involving the identification of a specific type of tumor-associated endothelial cells through single-cell data analysis, construction of an endothelial cell-related prognostic model, and validation of its performance (Fig. 1A). First, we subjected the scRNA data from nine samples, including three adjacent normal tissues and six PDAC tissues, selected from GSE212966, to quality control, resulting in a total of 47,731 cells. Subsequently, dimensionality reduction and clustering analyses were performed to obtain 38 clusters (Supplementary Fig. S1A and B). Based on reference cell markers and gene sets, we classified these clusters into 13 distinct cell types: epithelial cells, CD8+ T cells, CD4+ T cells, B cells, macrophages, endothelial cells, pericytes, fibroblasts, monocytes, dendritic cells, plasma cells, and mast cells (Fig. 1B). Subsequently, we analyzed gene expression within each cell cluster and observed that specific cell markers were highly expressed in their respective clusters (Fig. 1D). Notably, the endothelial cell cluster exhibited the expression of VWF, a marker associated with angiogenesis. Consistent expression patterns were observed for common markers, such as EPCAM, COL1A1, PECAM1, and PTPRC, representing epithelial cells, fibroblasts, endothelial cells, and immune cells, respectively (Supplementary Fig. S1E). We observed differences in the proportion of cell types between PDAC and adjacent tissues, except for endothelial cells, which were more abundant in PDAC (Fig. 1C). Furthermore, in the CRA001160 dataset, we analyzed 35 samples, including 24 PDAC tissues and 11 normal tissues, using dimensionality reduction and clustering analysis, resulting in the identification of 11 distinct cell clusters based on the previously mentioned cell subtypes (Supplementary Fig. S1C and D). We identified a significant number of endothelial cells in PDAC samples (Supplementary Fig. S2A). Therefore, we examined endothelial cells present in the PDAC microenvironment.

**Figure 1.**
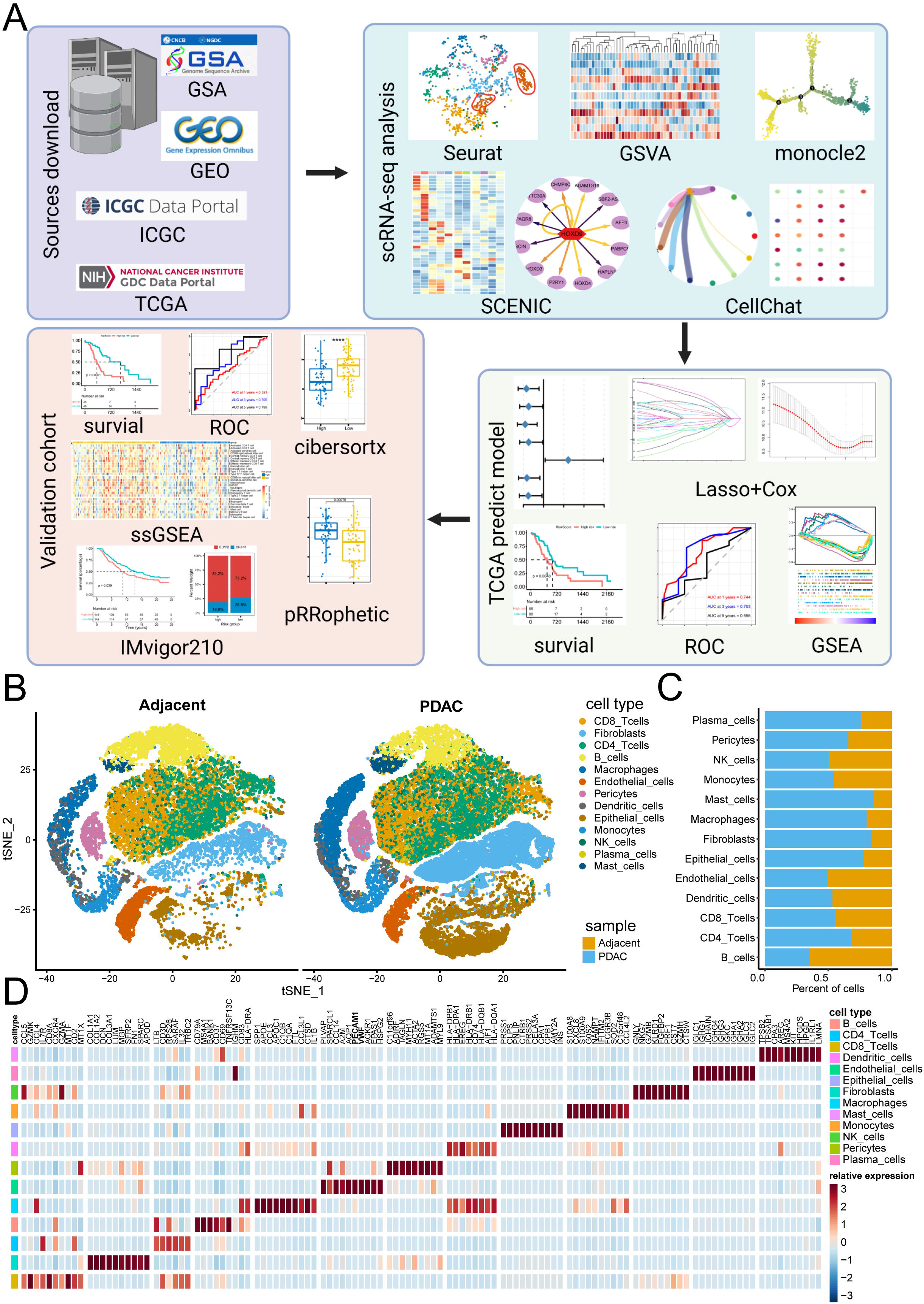
Single-cell Atlas of human adjacent pancreas and PDAC tissues. **A,** Workflow of this study. We downloaded publicly available data from PDAC patients and performed single-cell sequencing analysis to construct a prognostic model and validate its performance. **B,** TSNE plots of 16,228 cells from adjacent pancreas and 31,443 cells from tumor tissue of 6 PDAC patients, showing 13 clusters in each plot. Each cluster was shown in different colors. SCTransform was used to correct batch effects and constructed one TSNE based on all cells from adjacent tissue and tumor, and then split cells by these two tissues. **C,** Proportion of 13 major cell types in different sample are shown in bar plots. **D,** Heatmap shown the top DEGs (Wilcoxon test) in each 13 major cell types.

### 3.2 Tumor-associated COL4A1+ endothelial cells promote angiogenesis

Next, we performed dimensionality reduction and clustering on 2,008 endothelial cells from GSE212966, which were visualized using different colors (Fig. 2A). Among the 11 subclusters identified, endothelial cluster 6 showed a higher proportion exclusively in PDAC samples than in adjacent tissues (Fig. 2B). Notably, endothelial cells in cluster 6 exhibited elevated expression of COL4A1 and VWF (Fig. 2C). In the CRA001160 cohort, 8,547 endothelial cells were analyzed using dimensionality reduction and clustering techniques. We discovered a specific subcluster, endothelial cluster 1, which demonstrated the highest expression of COL4A1 and was predominantly present in tumor tissues (Supplementary Fig. S2B-D). Gene Set Variation Analysis (GSVA) performed on tumor hallmarks indicated that COL4A1+ endothelial cells were enriched in pathways associated with angiogenesis, E2F targets, G2M checkpoints, and mitotic spindles (Fig. 2D). Therefore, the presence of COL4A1+ endothelial cells signifies a distinct subpopulation of endothelial cells specifically associated with tumor neovascularization in PDAC.

**Figure 2.**
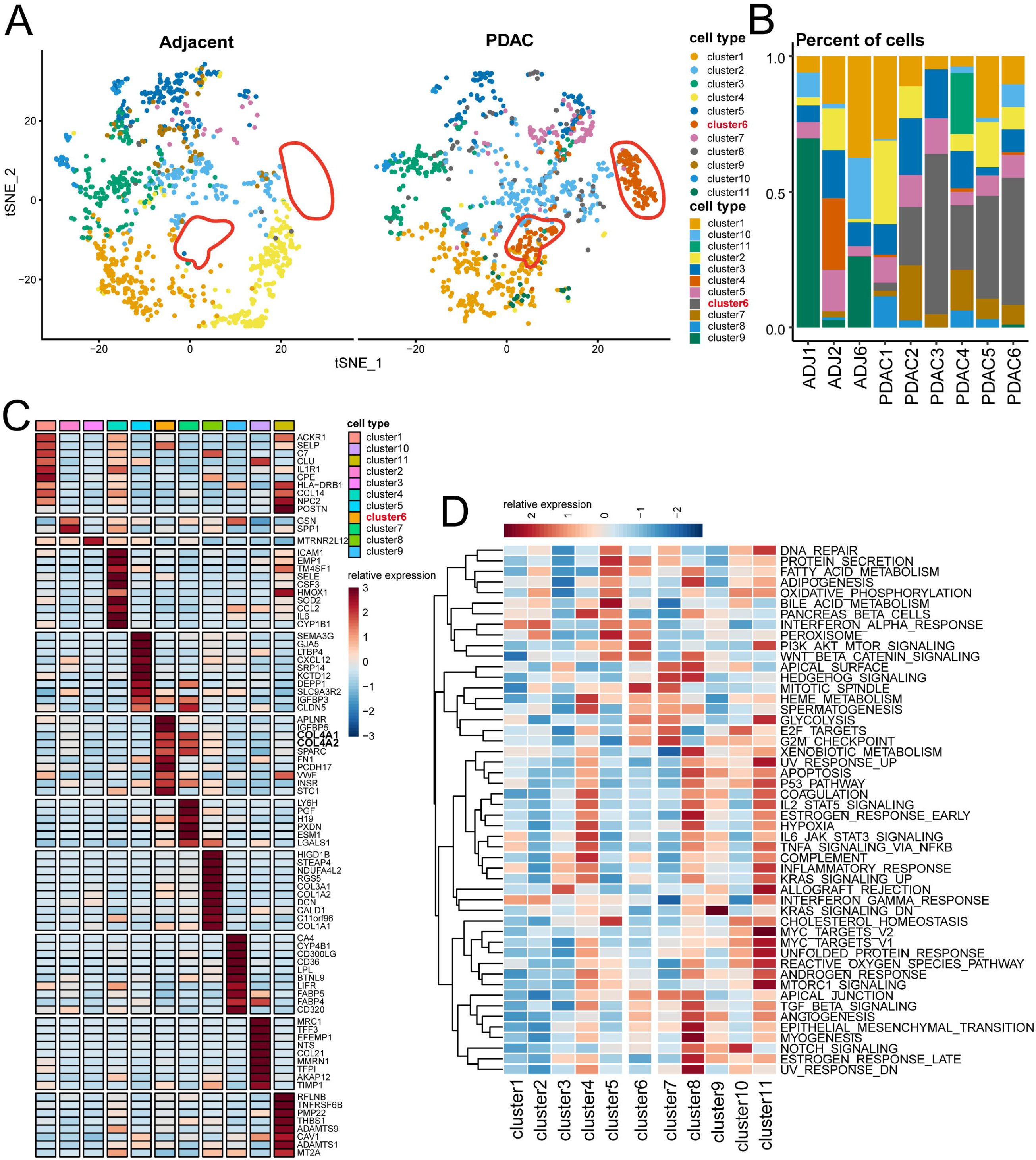
Characterization of cancer associated endothelial cells in PDAC. **A,** TSNE showing the composition of 2,008 endothelial cells colored by cluster. Red dashed circle shows COL4A1+ endothelial cells. **B,** Proportion of 12 endothelial subclusters showing in bar plots in different donors was shown. **C,** Heatmap to show the top DEGs (Wilcoxon test) in each 12 endothelial subclusters. **D,** Activation of hallmark pathways (scored per cell by GSVA function “ssgsea”) in 12 endothelial subclusters.

### 3.3 Transcription factor HOXD9 exhibited specific activation in COL4A1+ endothelial cells

To investigate the potential involvement of transcription factor regulation in angiogenesis, we employed the SCENIC software package to calculate the activity of transcription factor regulons and networks across 11 endothelial cell clusters. Notably, the transcription factors HOXD9, SOX4, and LEF1 were activated in COL4A1+ endothelial cells (Fig. 3A). Furthermore, by correlating the gene expression of transcription factors with specific cell types, we identified a specific association between HOXD9 and COL4A1+ endothelial cells (Fig. 3B). HOXD9 expression was observed in COL4A1+ endothelial cells in both cohorts (Fig. 3C and Supplementary Fig. S2E-F). In addition, the downstream targets regulated by HOXD9 encompass various genes involved in developmental regulation, including members of the HOXD family (HOXD3, HOXD4, and HOXD9). Other targets include CHMP4C, which stimulates growth factor receptors; AFF3, which promotes tumor formation; and SCIN, PAQR8, P2RY1, TTC30A, HAPLN1, and PABPC5, which are implicated in cellular maturation, transport, and metabolism (Fig. 3D). Moreover, the survival outcomes of PDAC patients correlated with the expression of HOXD9, whereby higher HOXD9 expression was associated with poorer prognosis in TCGA (p=3e-04) cohorts (Fig. 3E). Immunofluorescence staining demonstrated colocalization of HOXD9 and endothelial cells (CD31) in PDAC tissues (Fig. 3F). Thus, HOXD9 may specifically regulate neovascularization of COL4A1+ endothelial cells, potentially influencing unfavorable patient outcomes.

**Figure 3.**
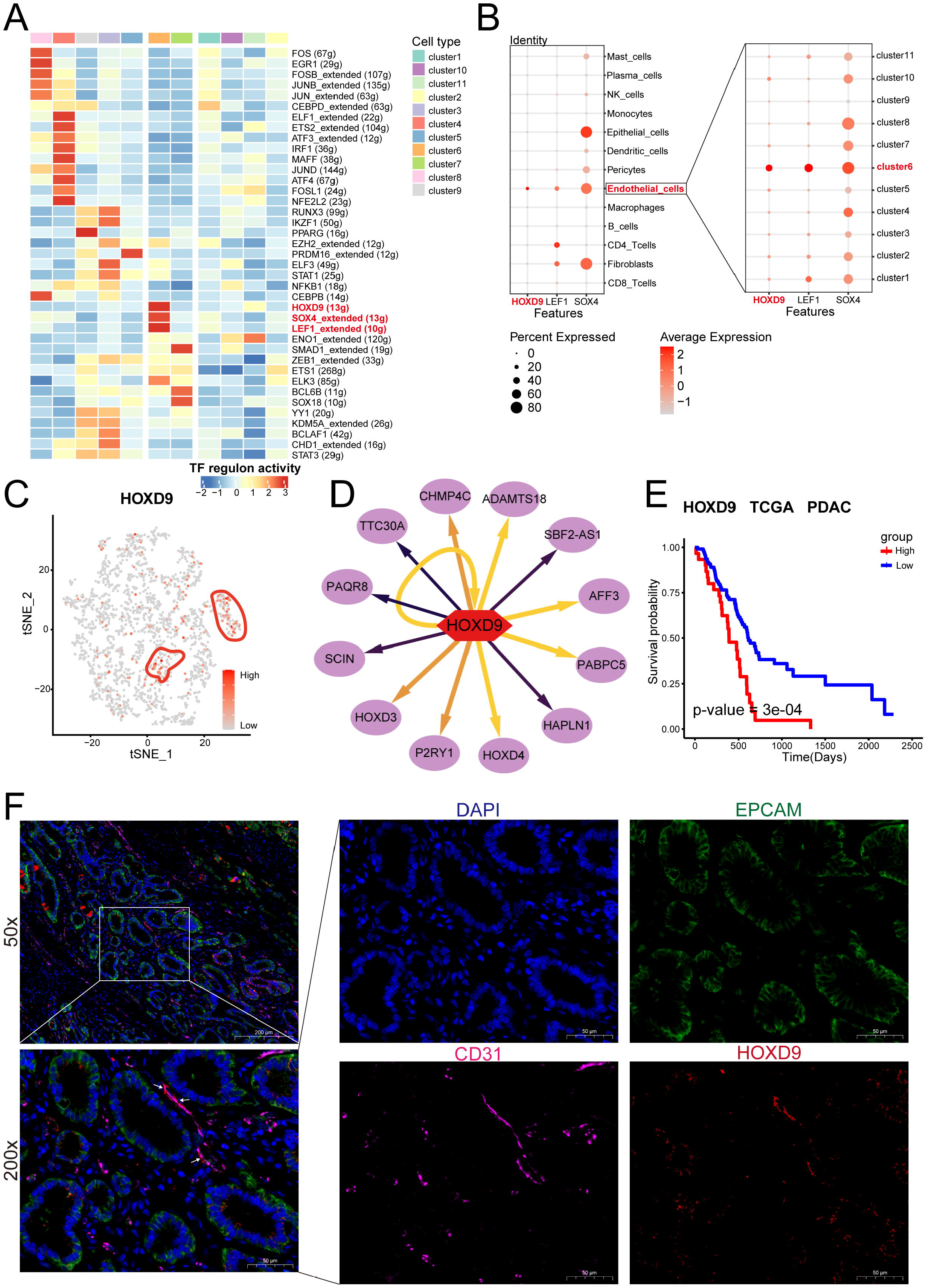
High expression of HOXD9 in COL4A1+endothelial cells associated with worse patient survival. **A,** Heatmap shows normalized activity of top TF regulons in endothelial subclusters predicted by R package SCENIC. **B,** Dotplot shows the relative expression (z-score) of three transcription factors (TFs) genes associated with COL4A1+endothelial cluster 6 in 13 major cell types (left) and 12 endothelial subclusters (right). **C,** TSNE plot showing expression of HOXD9. scale by z-score. **D,** The regulatory network of HOXD9 target genes. **E,** The Kaplan–Meier overall survival curves of PDAC patients in TCGA stratified by expression of HOXD9. **F,** Representative IF staining of human PDAC tissue. DAPI (blue), EPCAM (green), CD31 (pink), HOXD9 (red), in individual and merged channels are shown. Bar, 50 μm.

### 3.4 Developmental trajectory and stemness of COL4A1+ endothelial Cells in PDAC tumors

We constructed a developmental trajectory for endothelial cell differentiation to investigate the cellular state of COL4A1+ endothelial cells. Using CytoTRACE software, we predicted the cell stemness scores for the 11 endothelial cell clusters, revealing that cluster 8 exhibited the highest stemness, whereas cluster 9 had the lowest stemness (Fig. 4A, B). Moreover, COL4A1+ endothelial cluster 6 was in the mid-stage of differentiation (Fig. 4C). Furthermore, using Monocle2 software, we generated a developmental trajectory for endothelial cells, indicating a potential progression of endothelial differentiation from cluster 8 towards normal and neoplastic endothelial cell types (Fig. 4D). Additionally, the expression of HOXD9 aligned with the developmental trajectory of COL4A1+ endothelial cluster 6 (Fig. 4E, F). These findings suggest that COL4A1+ endothelial cells originate from a stem-like endothelial differentiation state and exhibit tumor-specific characteristics.

**Figure 4.**
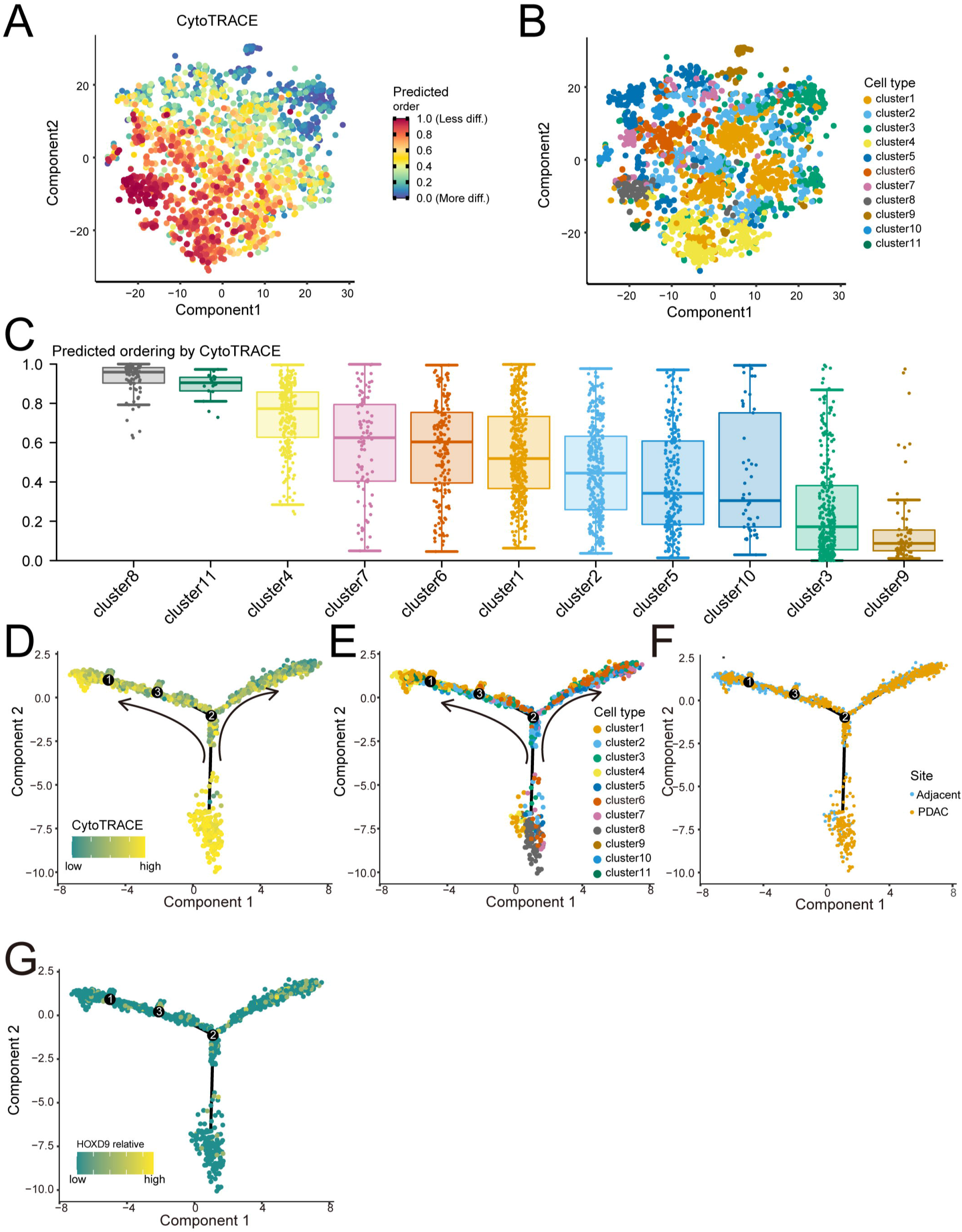
Differentiation and development of endothelial cells. **A**, Prediction of the differentiation and developmental status of endothelial cell clusters based on the cytoTRACE software. Dark-green indicates lower scores (low stemness) while dark-red indicates higher scores (high stemness). **B,** The cellular subpopulation states corresponding to **A**. Each cluster was shown in different color as Fig. 2A. **C,** Box plot shows the differentiation scores of the 11 endothelial cell clusters. **E,** UMAP plot shows the endothelial cell clusters within the developmental trajectory. **F,** UMAP plot shows the site of endothelial cell within the developmental trajectory. **G**, UMAP plot shows the expression pattern of the transcription factor HOXD9 within the developmental trajectory.

### 3.5 Epithelial cells, fibroblasts, and macrophages promote the generation of tumor vasculature in COL4A1 endothelial cells

We utilized the powerful signaling pathway analysis capability of CellChat to uncover the signaling interactions between COL4A1+ endothelial cells and other cell types. All signals were categorized into incoming and outgoing types, and we found that COL4A1+ endothelial cells exhibited a higher number of signals with malignant epithelial cells, fibroblasts, macrophages, dendritic cells, mast cells, monocytes, and pericytes (Supplementary Fig. S3A-B). Consequently, we visualized the specific signals between COL4A1+ endothelial cells and these seven cell types, including the VEGF, LAMININ, and COLLAGEN signaling pathways. The results showed that the incoming signals of COL4A1+ endothelial cells included the VEGF signaling pathway, and the outgoing signals included the LAMININ and COLLAGEN signaling pathways. The results revealed that COL4A1+ endothelial cells received incoming signals, such as the VEGF signaling pathway, and generated outgoing signals, including the LAMININ and COLLAGEN signaling pathways. Specifically, COL4A1+ endothelial cells promoted angiogenesis by receiving VEGFA and VEGFB signals secreted by dendritic cells, epithelial cells, fibroblasts, macrophages, mast cells, monocytes, and PGF signals secreted by pericytes through their own receptors VEGFR1/2 (Fig. 5A-C). LAMININ, a basement membrane protein, and COLLAGEN, an extracellular matrix protein, are crucial components of the tumor microenvironment. Aberrant secretion of LAMININ family proteins (LAMC1, LAMB1/2, and LAMA4/5) and COLLAGEN family proteins (COL4A1/2 and COL1A1/2) by COL4A1+ endothelial cells, epithelial cells, fibroblasts, and pericytes, along with the expression of integrin receptors (ITGA and ITGB) in the extracellular matrix, contributed to the modulation of the tumor microenvironment (Fig. 5D-G). Therefore, tumor-associated COL4A1+ endothelial cells promote tumor proliferation, and malignant cells facilitate neovascularization of COL4A1 endothelial cells.

**Figure 5.**
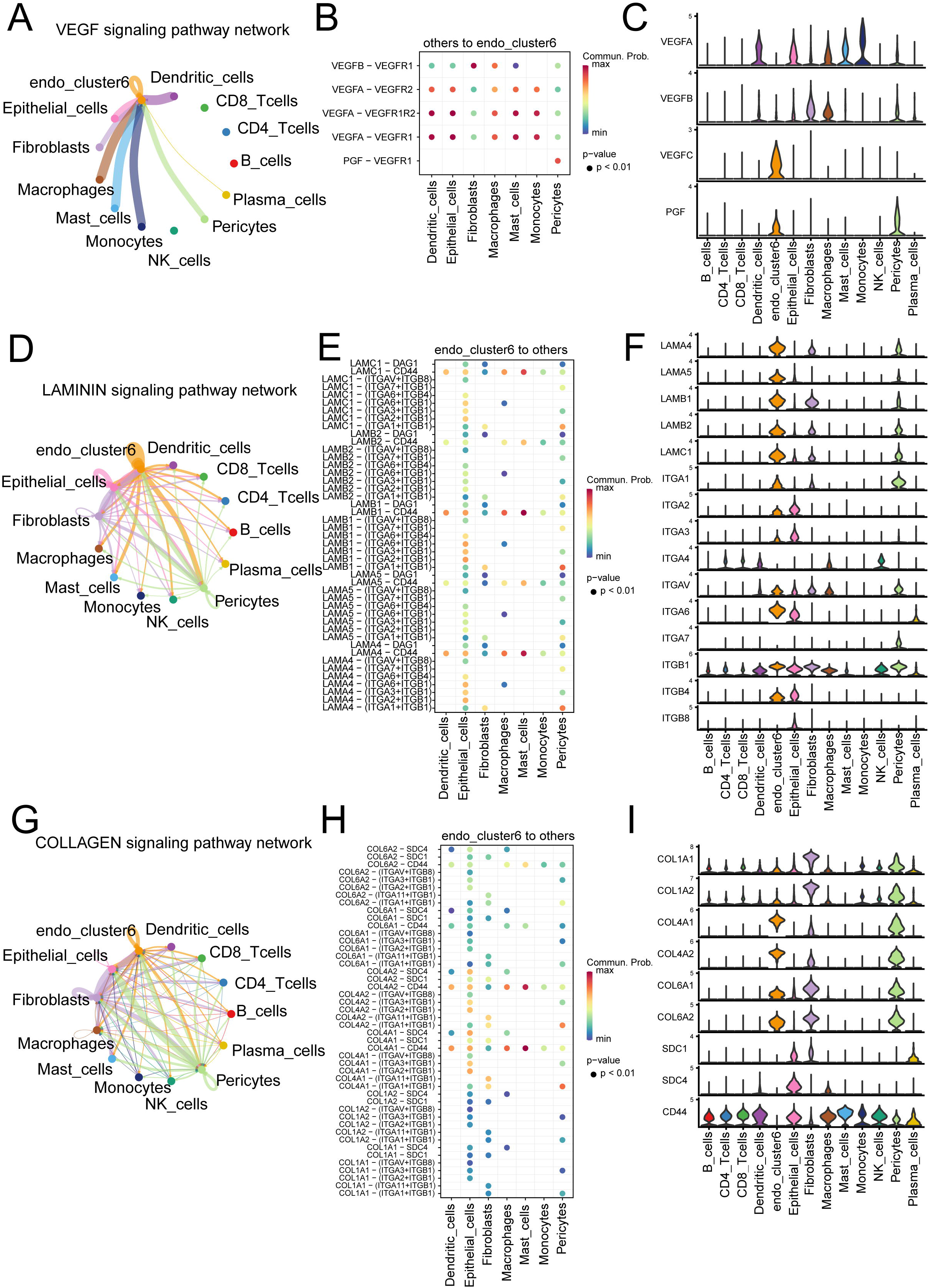
Intercellular interactions among COL4A1+ endothelial cluster 6 and malignant cells in PDAC. **A,** Cell–cell communications show VEGF signaling pathway network from other 12 major cell types to COL4A1+ endothelial cluster 6 by Cell chat analysis. **D, G,** Cell–cell communications show LAMININ signaling pathway network (**D**), signaling pathway network (**G**) from COL4A1+ endothelial cluster 6 to other 12 major cell types. **B, E, H,** Dot plots showed selected ligand-receptor interactions between COL4A1+ endothelial cluster 6 and other 7 major cell types. The ligand-receptor interactions and cell-cell interactions are indicated at columns and rows, respectively. The means of the average expression levels of two interacting molecules are indicated by colour heatmap (right panel), with blue to red representing low to high expression. The P-values were indicated by circle size in one-sided permutation test. **C, F, I,** Violin plots to show expression levels of multiple legends and receptors in VEGF signaling pathway (**C**), LAMININ signaling pathway (**F**), and COLLAGEN signaling pathway (**I**) in different cell types.

### 3.6 Construction and validation of the prognostic risk model TaEMS

Next, we investigated the prognostic value of COL4A1+ endothelial cells in patients with PDAC. We identified 118 differentially expressed genes (DEGs) that were the intersection of COL4A1+ endothelial DEGs, 550 DEGs from GSE212966, and 217 DEGs from CRA001160 (Fig. 6A). These DEGs were subjected to univariate Cox regression analysis (Supplementary Fig. S4B). From the univariate Cox regression analysis, seven genes with significant p values were selected and used to construct the LASSO regression model (Fig. 6B). Through ten-fold cross-validation, five genes with non-zero coefficients, namely DDIT4, TIE1, SEMA6B, PLCB1, and LYZ, were identified as the optimal model (Fig. 6C-D and Supplementary Fig. S4A). TaEMS exhibited good stratification performance for overall survival (OS) in PDAC patients based on TCGA database (p=0.0048) (Fig. 6E). Furthermore, TaEMS showed favorable prognostic significance for OS in two independent cohorts, the ICGC (p<0.0001) and GEO databases (p=0.0095) (Fig. 6F-G). Subsequently, we calculated the area under the curve (AUC) for the 1-, 3-, and 5-year survival rate predictions using the TaEMS model. The ROC curve analysis yielded AUC values of 0.744, 0.753, and 0.595 for 1-, 3-, and 5-year survival rates, respectively, in the TCGA training set (Fig. 5H); 0.755, 0.65, and *NA* in the ICGC validation set (Fig. 6I); and 0.591, 0.705, and 0.799 in the GEO validation set (Fig. 6J). These results indicate the potential predictive ability of TaEMS for short-term survival rates in patients. Moreover, the high-risk group showed enrichment in cell cycle and DNA replication pathways, whereas the low-risk group exhibited enrichment in the T cell receptor signaling pathway, chemokine signaling pathway, and cytokine-cytokine receptor interaction (Supplementary Fig. S4C). These findings were consistent with the enrichment results of COL4A1+ endothelial cells from the scRNA-seq analysis of mentioned before. Thus, TaEMS demonstrated predictive value for patient prognosis and survival.

**Figure 6.**
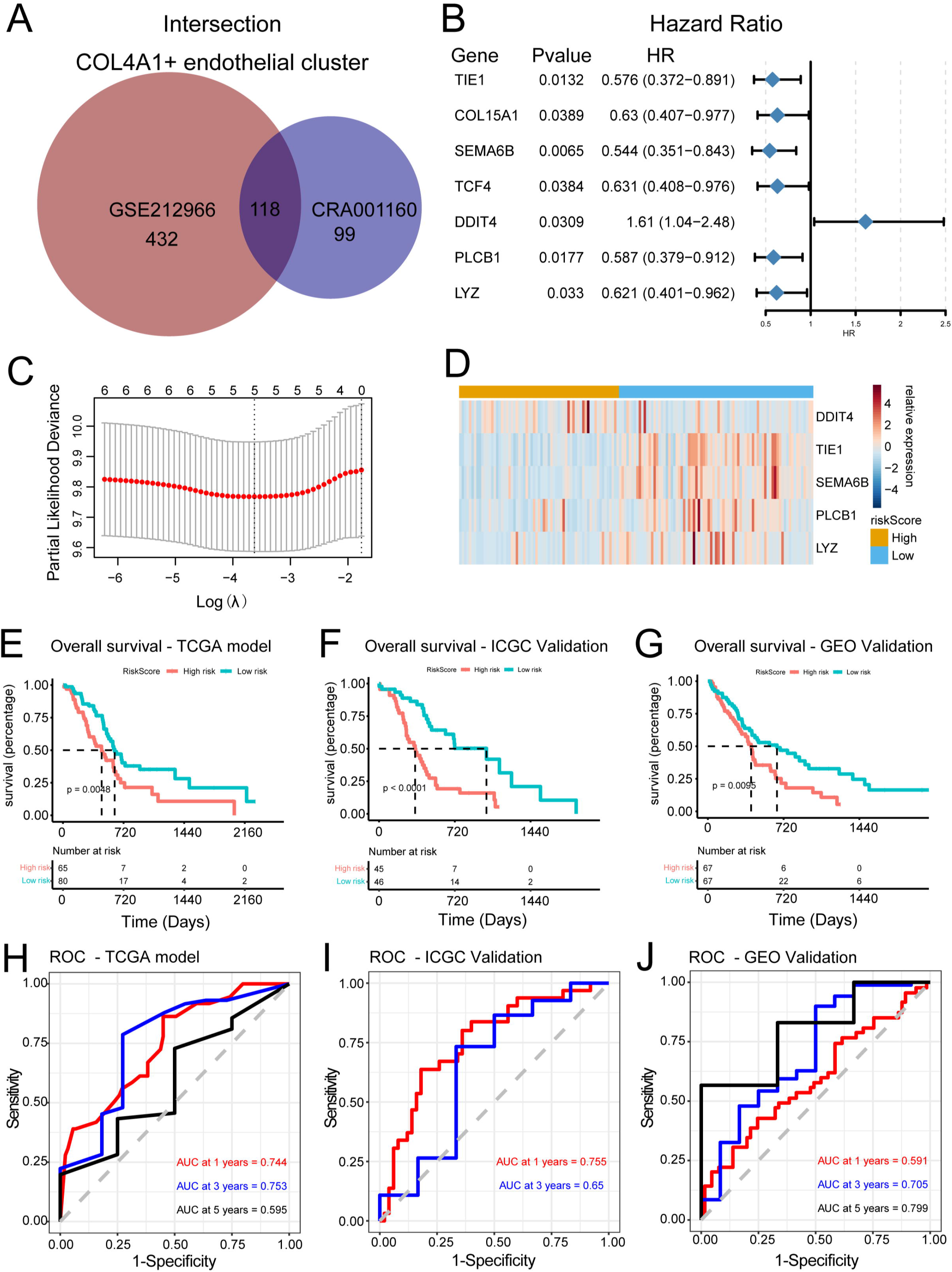
Establishment and validation of TaEMS in the TCGA PDAC and External cohort. **A,** Circle plot shown the intersection 118 DEGs of the TaEMS. **B**, Forest plot of the 7 most significant prognostic genes associated with PDAC prognosis. HR is the hazard ratio and 95% CI is the 95% confidence interval. **C**, The plot of the determination of the tuning parameter lambda in the LASSO regression model. The X axis is the log (lambda) and Y axis is the partial likelihood deviation value. The lambda value corresponding to the smallest value is the best. **D,** The expression heat map of the 5 TaEMS selected from the TCGA PDAC dataset. **E-G,** Kaplan–Meier survival curve in the TCGA model (**E**), and validation of ICGC dataset (**F)**, GEO dataset (**G**). The X axis represents time, while the Y axis represents survival. Different colours represent different groups. The p-value is based on the log-rank test. **H-J,** ROC curves for predicting the risk of death at 1, 3, and 5 years in the TCGA model (**H**), ICGC dataset (**I**), GEO dataset (**J**) based on group by TaEMS risk score.

### 3.7 TaEMS predicted immune cell infiltration and the potential benefit of antiangiogenic therapy in PDAC

Due to the adverse effects of angiogenesis on tumor immunity, we further investigated the relationship between TaEMS and immune cell infiltration in patients with PDAC. Gene sets for 28 immune cell types were obtained from previous studies[28], and enrichment analysis using ssGSEA was performed on immune cells from 145 patients with PDAC from TCGA. Samples were divided into two groups based on their high- and low-risk scores. We found that patients in the low-risk group had higher enrichment scores of immune cells than those in the high-risk group (Fig. 7A). Additionally, using ESTIMATE software, we observed that the high-risk group had lower immune, stromal, and ESTIMATE scores, and higher tumor purity than the low-risk group (Fig. 7B), indicating a potential negative correlation between TaEMS risk scores and immune cell infiltration levels. Furthermore, levels of immune checkpoints such as PD1, PD-L1, and CTLA4 were found to be higher in the low-risk group than in the high-risk group (Supplementary Fig. S5G), suggesting a potential association between TaEMS and immunotherapy. Subsequently, the performance of TaEMS was tested in an immunotherapy cohort, IMvigor210, and was found to be prognostically significant (p=0.039) in this cohort (Supplementary Fig. S5E). The tumor mutational burden (TMB) between the high-risk and low-risk groups of PDAC patients was similar, with KRAS, TP53, and SMAD4 mutations being the main alterations (Supplementary Fig. S6A). A similar trend was observed in the IMvigor210 cohort (Supplementary Fig. S5C). However, patients in the low-risk group had a higher tumor neoantigen burden (TNB) than those in the high-risk group (Supplementary Fig. S5D). Moreover, the low-risk group had an 8% higher rate of immunotherapy response (complete response/partial response, CR/PR) than the high-risk group (Supplementary Fig. S5F). Although this cohort was not specific to PDAC immunotherapy, TaEMS could serve as a useful biomarker for tumor immunity to determine which patients may benefit from immunotherapy. Owing to the significance of TaEMS in relation to TAECs and angiogenesis, we further investigated the role of TaEMS in antiangiogenic therapies. The pRRophetic tool was used to calculate the IC50 values of 138 drugs in 145 PDAC tumor samples. The samples were categorized into high and low-risk groups, and notable differences in drug sensitivity were observed among the groups for 40 drugs, including five antiangiogenic agents (Fig. 7C and Supplementary Tables S3). These antiangiogenic drugs included ponatinib (AP24534) (p=0.00047), AZ628 (p=0.00044), GSK269962A (p=0.0021), pazopanib (p=0.00076), and sunitinib (p=0.016), with low-risk patients exhibiting lower IC50 values. Collectively, low-risk scores were positively correlated with immune cell infiltration and higher checkpoint expression, suggesting that low-risk patients may potentially benefit from both immunotherapy and antiangiogenic drugs.

**Figure 7.**
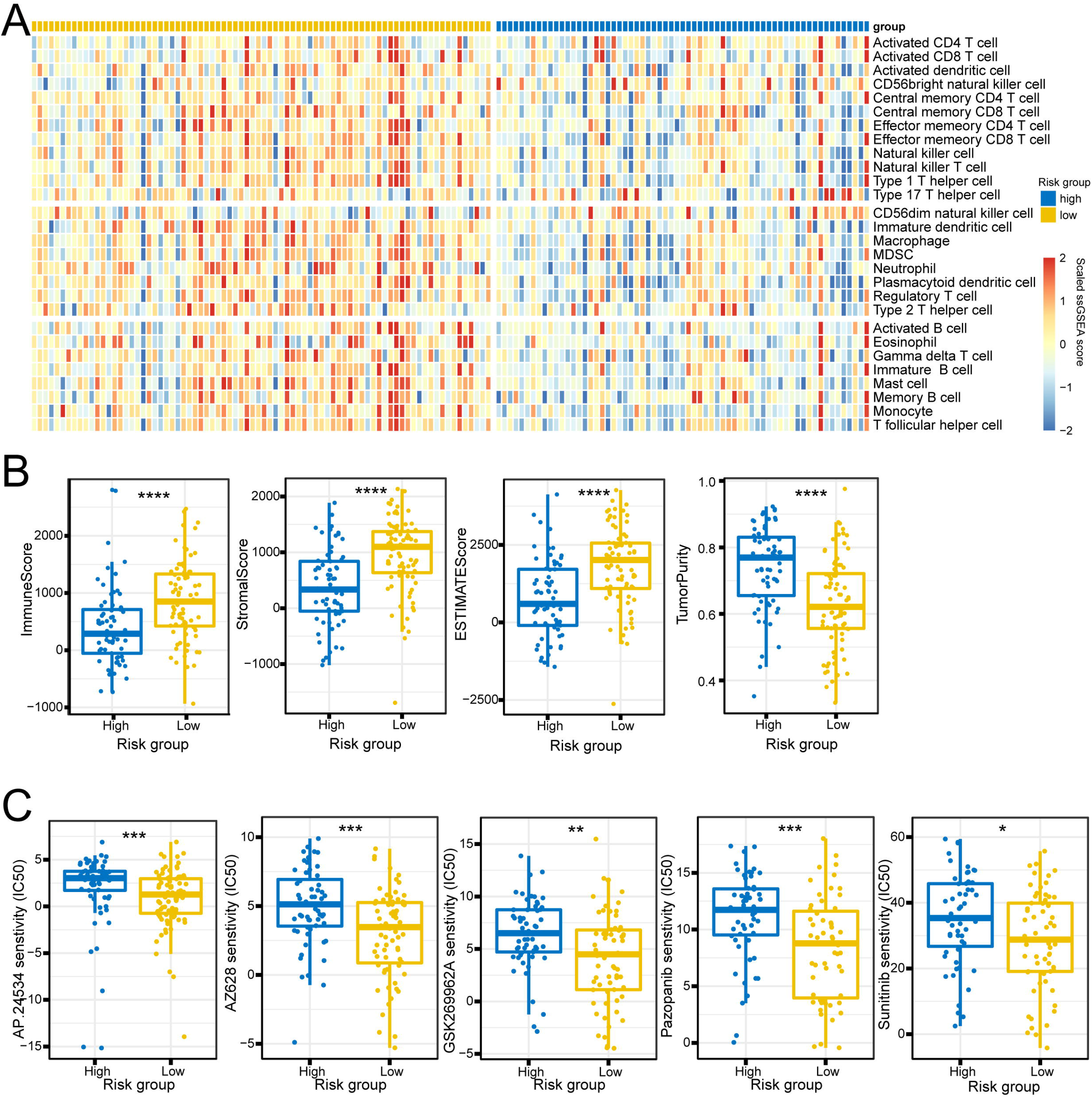
The TaEMS associated with infiltration of immune cells in TME and Drug sensitivity. **A**, Single-sample gene set enrichment analysis identifying the relative infiltration of immune cell populations for 145 tumor samples with available RNA-seq data in TCGA. **B**, Differences among immune score, stromal score, ESTIMATE score and tumor purity between high-risk and low-risk groups. **C,** The Comparison of drug sensitivity (IC50) in AP.24534, AZ628, GSK269962A, Pazopanib, and Sunitinib between high-risk and low-risk groups. Statistics based on Wilcoxon test. Statistics based on Wilcoxon test. P< 0.05*, <0.01**, <0.001***, <0.0001 ****.

## 4. Discussion

With the advancement of single-cell sequencing technologies, researchers have gained the ability to delve deeper into the molecular characteristics of the tumor immune microenvironment. However, despite their critical involvement in angiogenesis and potential impact on prognosis and treatment response, the role of TAECs has not received adequate attention in recent studies. The association between TAECs and patient prognosis has been demonstrated in various solid tumors[29–31]. In a study by Sun et al., the emerging VEGF signaling mediated by PGF-VEGFR1 between cancer-associated fibroblasts (CAFs) and endothelial cells in intrahepatic cholangiocarcinoma (ICC) highlighted the potential of targeting CAFs to improve immune checkpoint blockade (ICB) therapy[29]. Daum et al. reported that antiangiogenic therapies could benefit highly vascularized tumors, such as non-small cell lung cancer [30]. Furthermore, Xie et al. utilized single-cell sequencing to uncover abnormal permeability of tumor endothelial cells in glioblastoma multiforme (GBM), offering novel insights for designing rational treatment strategies[31]. Motivated by these studies, our investigation focused on TAECs in the context of PDAC, using marker genes identified through scRNA-seq analysis. We identified a subset of COL4A1+ endothelial cells that displayed a close association with tumor cells and protumor cells within the tumor microenvironment.

In this study, we discovered a subset of endothelial cells expressing high levels of COL4A1 in two independent cohorts. Notably, these cells were exclusively found within the tumor region of PDAC and not in adjacent nontumor or normal tissues. We designated these COL4A1+ endothelial cells as TAECs. COL4A1 belongs to the collagen family, which is the most abundant protein in the extracellular matrix and is a vital component of the tumor microenvironment. Our research revealed the interactions of COL4A1+ endothelial cells with various cell types, including tumor cells, tumor-associated fibroblasts, macrophages, and pericytes, through the COLLAGEN signaling pathway. Specifically, we identified key signaling pathways involving COL4A1-ITGA/B, COL4A1-CD44, and COL4A1-SDC1/4. In line with our findings, Wang et al. discovered that COL4A1 is highly expressed in hepatocellular carcinoma (HCC) cells and that its upregulation promotes HCC cell proliferation, migration, and invasion [32]. Similarly, Cui et al. reported that silencing COL4A1 in gastric cancer (GC) cells inhibited cell viability, migration, and invasion[33]. These studies provide further support for our findings and indicate that in PDAC, tumor cells themselves either do not express COL4A1 or express it at low levels. Instead, COL4A1+ endothelial cells stimulate the proliferation, migration, and invasion abilities of tumor cells through the COLLAGEN signaling pathway.

Furthermore, our single-cell data analysis revealed a distinct activation of the HOXD9 transcription factor in COL4A1+ endothelial cells. These cells exhibited associations with critical biological processes, including cell cycle regulation, DNA repair, E2F targets, G2M checkpoints, protein secretion, and mitotic spindle processes. HOXD9 belongs to the HOX gene family, which plays a vital role in cellular development and regulates processes such as proliferation, apoptosis, and cell migration. Previous studies have provided insights into the role of HOXD9 in cancer progression. Lv et al. demonstrated that HOXD9 is highly expressed in invasive hepatocellular carcinoma (HCC) and promotes HCC cell migration, invasion, and metastasis [34]. Similarly, Zhu et al. reported that HOXD9 transcriptionally activated RUFY3, leading to enhanced proliferation and invasion of gastric cancer (GC) cells[35]. However, our findings present a different perspective, as tumor cells themselves do not express HOXD9. Instead, the activation of HOXD9 in COL4A1+ endothelial cells may contribute to tumor angiogenesis, facilitating the proliferation or metastasis of PDAC cells. Based on our scRNA-seq analysis, we propose a plausible mechanism by which tumor cells release VEGFA/B, stimulating abnormal proliferation of endothelial cells through the VEGF signaling pathway. This in turn activates HOXD9 in endothelial cells, regulating their proliferation and migration, ultimately leading to angiogenesis. Additionally, TAECs release molecules involved in the COLLAGEN and LAMININ signaling pathways into the extracellular matrix, further promoting the proliferation and migration of tumor cells.

Following our study, we developed a prognostic risk model called TaEMS, which demonstrated a strong predictive capability for PDAC patient prognosis. TaEMS comprises five marker genes associated with tumor-associated endothelial cells (DDIT4, TIE1, SEMA6B, PLCB1, and LYZ), all of which have implications for PDAC prognosis or endothelial cell proliferation. DDIT4, also known as DNA damage-inducible transcript 4, regulates cell growth, proliferation, and survival by inhibiting mTORC1 activity. Martha et al. previously reported that DDIT4 is associated with angiogenesis, and that genetic variations in DDIT4 contribute to breast cancer progression[36]. Additionally, Wenes et al. demonstrated that DDIT4-mediated mTOR inhibition leads to abnormal blood vessel formation [37]. TIE1 is a tyrosine kinase receptor, and Silvia et al. showed that conditional knockout of TIE1 in endothelial cells reduced tumor angiogenesis in mice and prolonged overall survival[38]. SEMA6B, a member of the semaphorin protein family, acts as a ligand for Plexin-A4. Plexin-A4 forms stable complexes with VEGFR2 receptors, enhancing VEGF-induced endothelial cell proliferation. Knockdown of SEMA6B leads to the silencing of Plexin-A4 and inhibits tumor formation in GBMU87MG cells[39]. PLCB1, also known as phospholipase C beta 1, plays a role in controlling neovascularization of endothelial cells upon activation. Wang et al. found that TRAX chelation enhances angiogenesis in the tumor microenvironment[40]. Consistent with our study, PLCB1 expression was lower in the high-risk group, potentially promoting tumor angiogenesis. As reported by Cui et al., it is involved in inflammatory responses and promotes leukocyte adhesion to endothelial cells[41]. Collectively, the identification of these marker genes in TaEMS provides potential targets for further laboratory experiments aimed at elucidating the molecular mechanisms underlying PDAC.

This study demonstrated the powerful predictive capability of TaEMS as a prognostic tool for patient prognosis in both training and validation cohorts. To further investigate the underlying mechanisms, we performed GSEA to identify the pathways associated with TaEMS. The high-risk group was enriched in the cell cycle pathway, whereas the low-risk group showed enrichment in pathways related to focal adhesion, vascular smooth muscle contraction, and immune-related processes. Dysregulated cell cycle control, which is closely associated with tumor proliferation and progression, may contribute to the adverse prognosis observed in high-risk group[42]. This finding aligns with our scRNA-seq analysis results, where COL4A1+ endothelial cells were associated with cell cycle and mitotic spindle cancer hallmarks. Additionally, considering the important role of tumor-infiltrating immune cells in tumor development[43], we compared immune cell infiltration between high-risk and low-risk groups using the ESTIMATE and ssGSEA algorithms. The results revealed lower immune cell infiltration, particularly of activated CD8+ T cells and NK cells, in the high-risk group. This suggests that tumors with high-risk scores exhibit characteristics of “cold tumors”[44] with reduced antitumor activity. Although suitable PDAC immunotherapy cohorts were not found, we also observed a higher immunotherapy response rate in low-risk patients than in high-risk score patients in the case of BLCA. This may also partially explain the lower survival rates observed in high-risk PDAC patients.

Furthermore, our study revealed significant differences in the sensitivity of TaEMS to antiangiogenic drugs (ponatinib, AZ628, GSK269962A, pazopanib, and sunitinib) in PDAC. Ponatinib is a kinase inhibitor targeting VEGFR, PDGFR, FGFR, and EGFR [45]. Ponatinib has been reported to inhibit PDGF, involved in angiogenesis within the tumor stroma[46]. AZ628 is a RAF kinase inhibitor that suppresses MITF and FRA-1, transcription factors associated with EGFR expression, leading to inhibited cell growth and migration in V600 BRAF wild-type melanoma cells[47]. GSK269962A, a ROCK1 inhibitor, induces apoptosis in acute myeloid leukemia cells by blocking the ROCK1/c-RAF/ERK pathway[48]. Pazopanib, initially developed as a small-molecule inhibitor of vascular endothelial growth factor receptors, is a tyrosine kinase inhibitor[49]. Lee et al. found that pazopanib improves progression-free survival in patients with advanced soft tissue sarcoma by inhibiting angiogenesis[50]. Sunitinib is a selective inhibitor of multiple receptor tyrosine kinases associated with tumor growth and angiogenesis[51, 52]. Early retrospective analyses by Deeks et al. confirmed the clinical efficacy of sunitinib in advanced gastrointestinal stromal tumors and advanced renal cell carcinoma with good patient tolerability[51]. These reports further support our findings that TaEMS may serve as a potential biomarker for selective treatment with antiangiogenic drugs, leading to improved therapeutic outcomes in patients with PDAC.

Despite these valuable findings, this study had several limitations. First, our investigation focused solely on endothelial cells in patients with PDAC, whereas the tumor immune microenvironment exhibited high heterogeneity. Therefore, further research on other tumor types is required. Second, TaEMS genes require further exploration at the protein level to understand their effects and prognostic significance. In addition, the limited predictive ability of TaEMS warrants further improvement. Last, the mechanistic analyses in our study were descriptive, and further research is needed to elucidate the underlying mechanisms linking tumor-associated endothelial cells, TaEMS, PDAC prognosis, and treatment.

In summary, this study provides valuable insights into the complex interactions between TAECs and tumor cells in PDAC. The activation of HOXD9 in COL4A1+ endothelial cells and its communication with tumor cells through the COLLAGEN signaling pathway have shed light on the mechanisms underlying tumor angiogenesis, and the promotion of tumor cell proliferation and migration. Furthermore, the development of TaEMS, a prognostic risk model based on five tumor-associated endothelial cell marker genes, has demonstrated strong predictive performance for PDAC patient prognosis and sensitivity to antiangiogenic therapy in solid tumors. TaEMS has the potential to serve as a prognostic biomarker, enabling personalized clinical decisions and identifying patients who may benefit from antiangiogenic treatments. These findings contribute to a deeper understanding of the tumor microenvironment and offer opportunities for improved management and treatment strategies for PDAC.

## Author contributions

JW and XDM conceived of the project. JW and YL performed the bioinformatics analysis under the supervision of XDM. QF and ZC performed multiple immunostaining experiments and conducted the image analysis. JW wrote the manuscript, and XL and XDM revised it. All authors have reviewed the manuscript and approved its submission and publication.

## Availability of data and materials

The bulk RNA-sequencing data for this study were derived from TCGA, GEO (accession number: GSE183795), and ICGC databases. The accession numbers for the single-cell RNA-sequencing data reported in this study were GSE212966 and CRA001160. All other relevant data supporting the key findings of this study are available within the article and its Supplementary Information file. For additional reasonable request, please feel free to contact the corresponding author.

## Code availability

The source code for this study is not provided. Interested researchers can contact the authors for more information or potential collaboration. Despite the code not being publicly available, we prioritize transparency and scientific collaboration within the research community.

## Declarations

### Ethics approval and consent to participate

Not applicable

### Consent for publication

All authors agree with the publication of this manuscript. Ethical approval and consent to participate. The data for this study were obtained from public databases, and no additional ethical approval was required.

PDAC: Pancreatic ductal adenocarcinoma
scRNA-seq: single-cell RNA sequencing
TAECs: Tumor-Associated Endothelial Cells
TaEMS: Tumor-associated Endothelial Marker Signature
GEO: Gene Expression Omnibus
GSA: Genome Sequence Archive
TCGA: The Cancer Genome Atlas
ICGC: International Cancer Genome Consortium
PCA: Principal component analysis
tSNE: t-distributed stochastic neighbor embedding
GSEA: Gene set enrichment analysis
VEGFR: Vascular Endothelial Growth Factor Receptor
PDGFR: Platelet-Derived Growth Factor Receptor
FGFR: Fibroblast Growth Factor Receptor
EGFR: Epidermal Growth Factor Receptor

## Acknowledgements

We thank all members of the laboratory for their experimental assistance.

