## Supplementary figures and images for "Characterization of Tumor-Associated Endothelial Cells and the Development of a Prognostic Model in Pancreatic Ductal Adenocarcinoma"

### Supplementary Figure 1

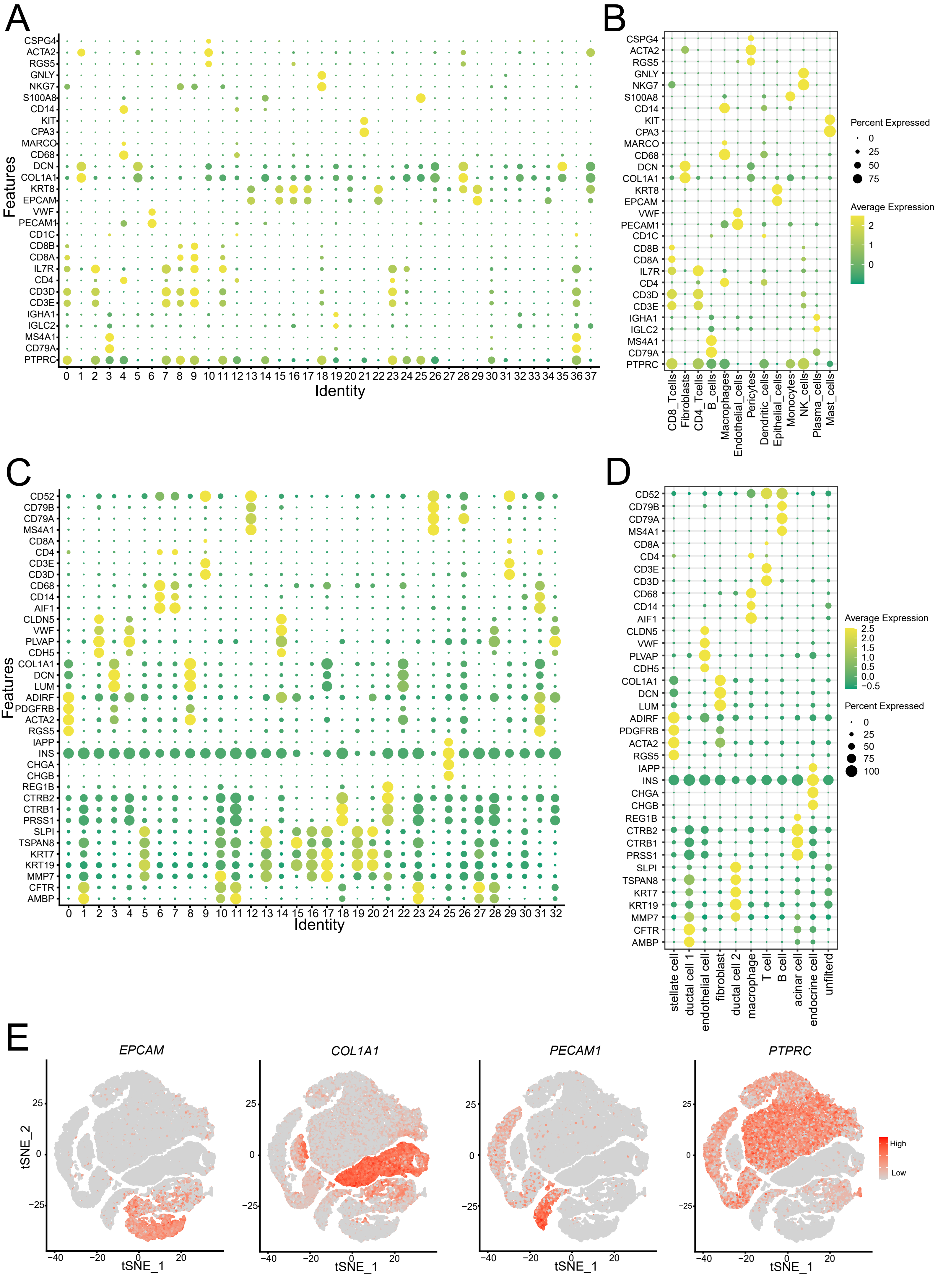

### Supplementary Figure 2

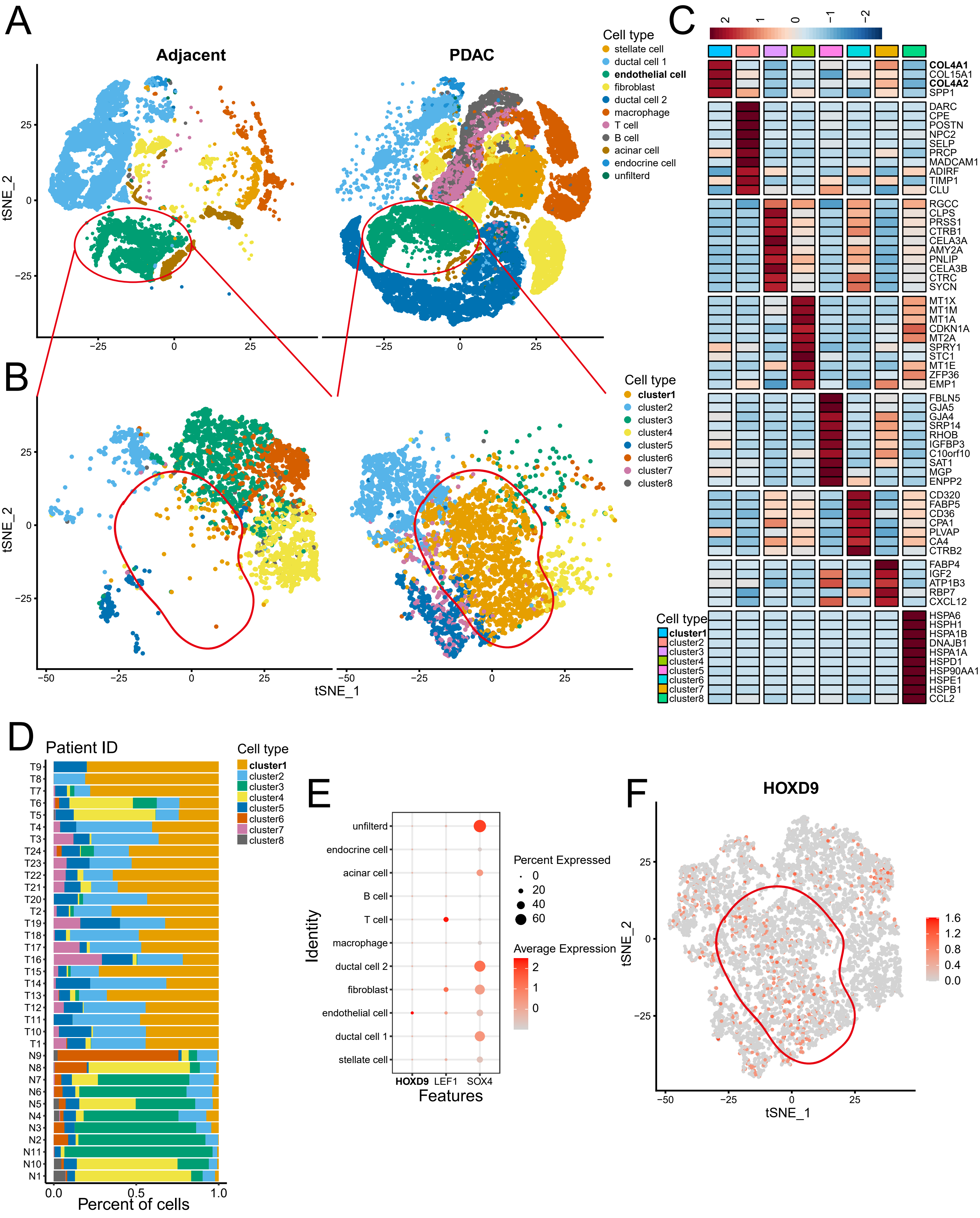

### Supplementary Figure 3

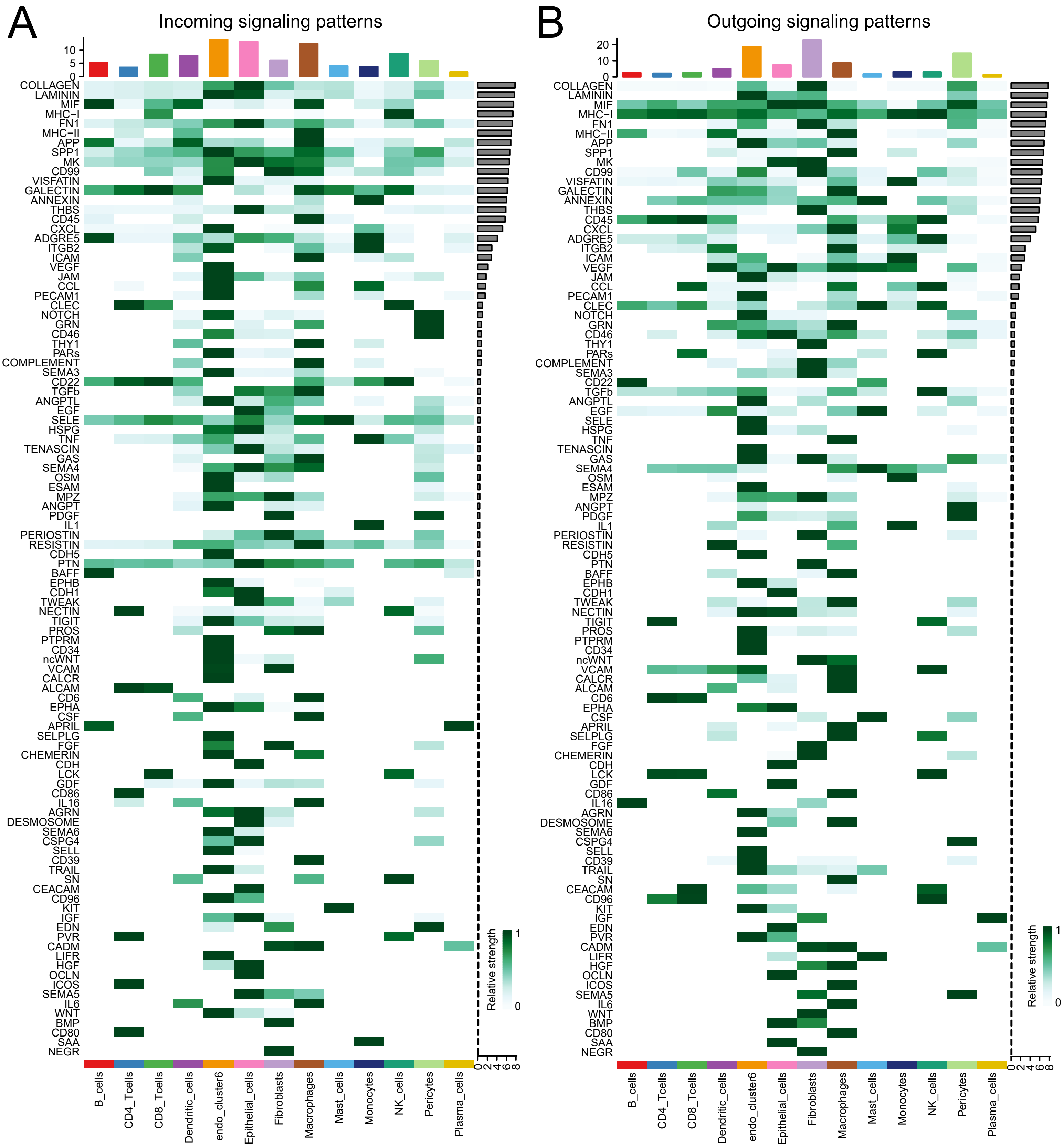

### Supplementary Figure 4

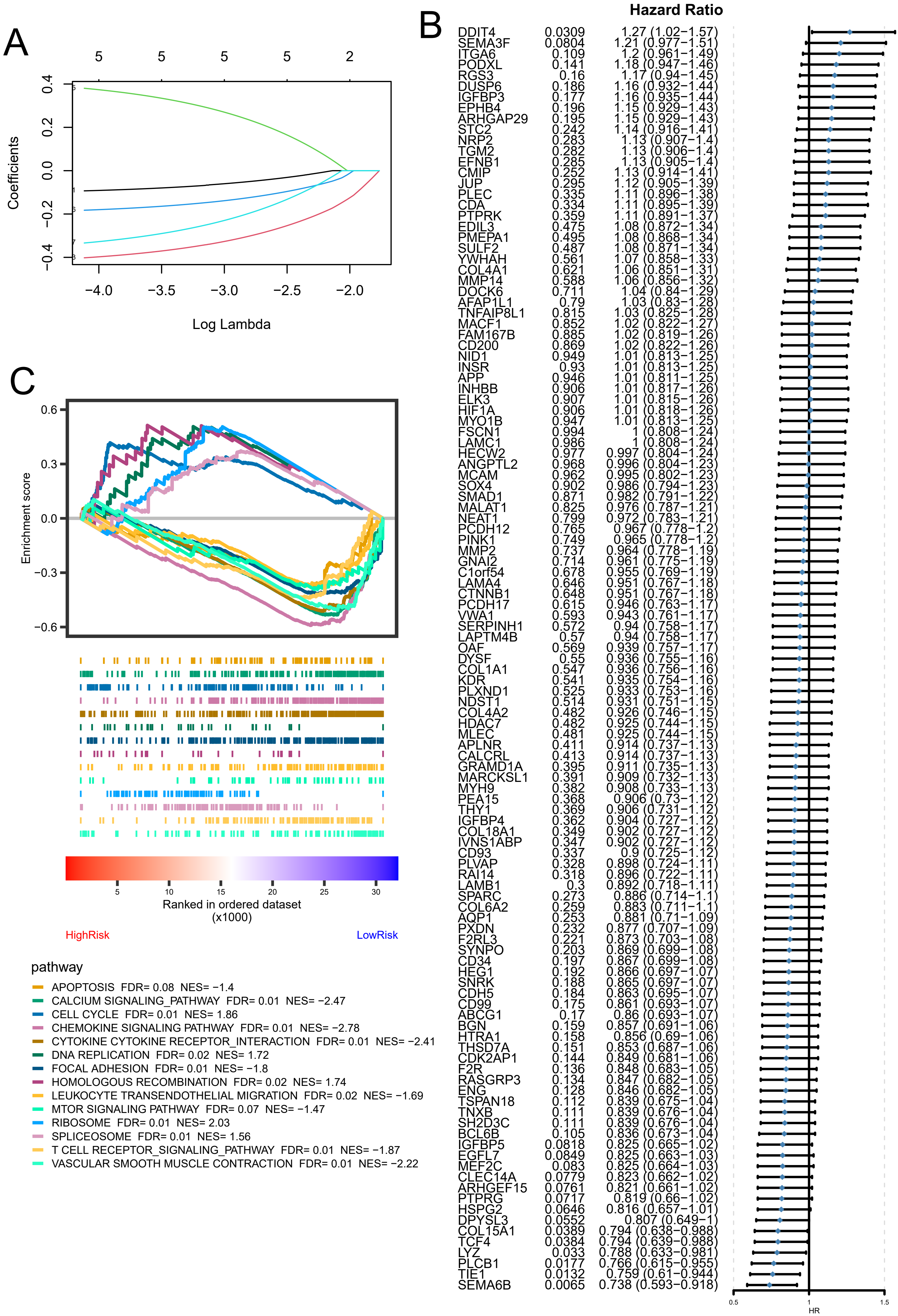

### Supplementary Figure 5

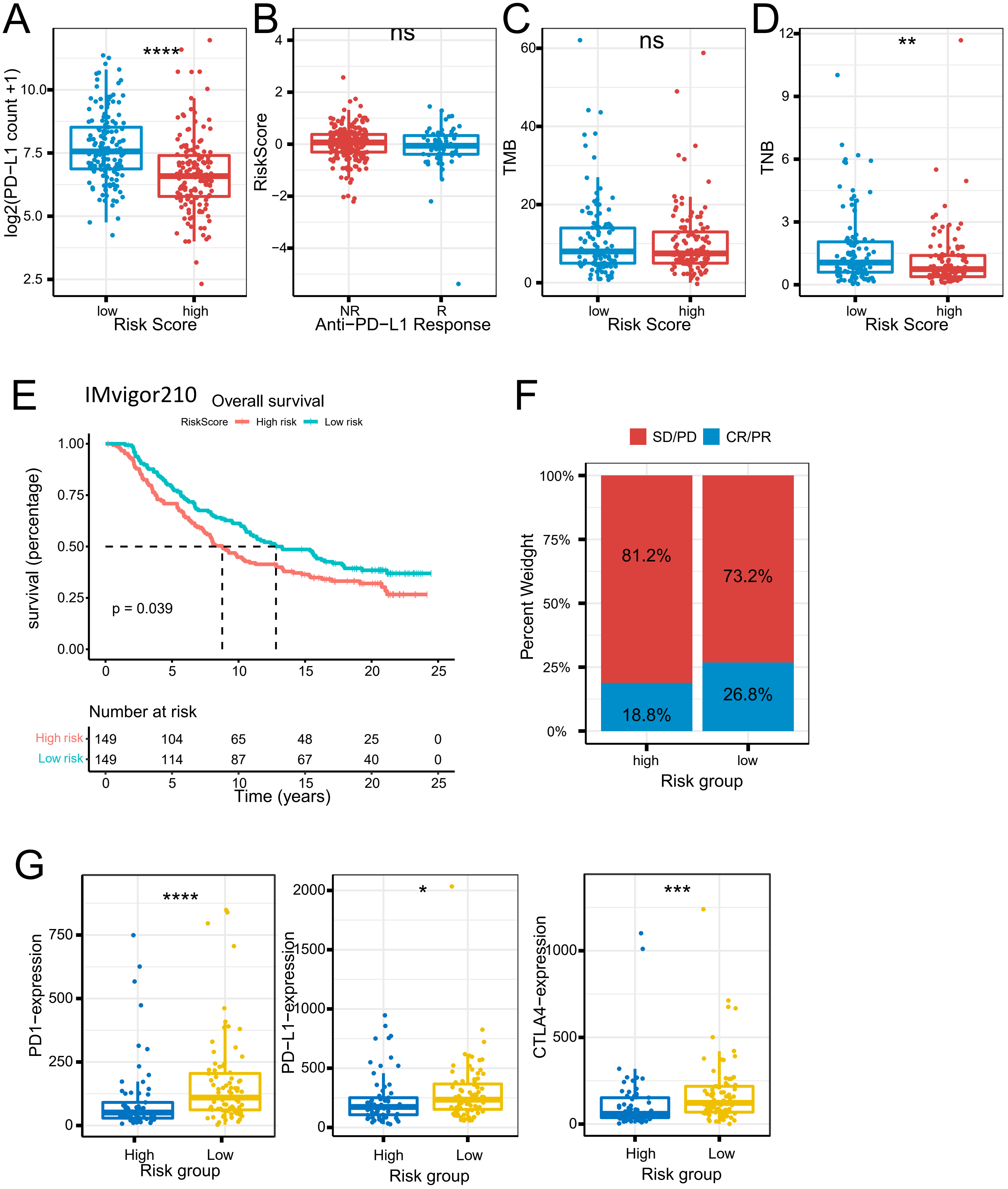

### Supplementary Figure 6

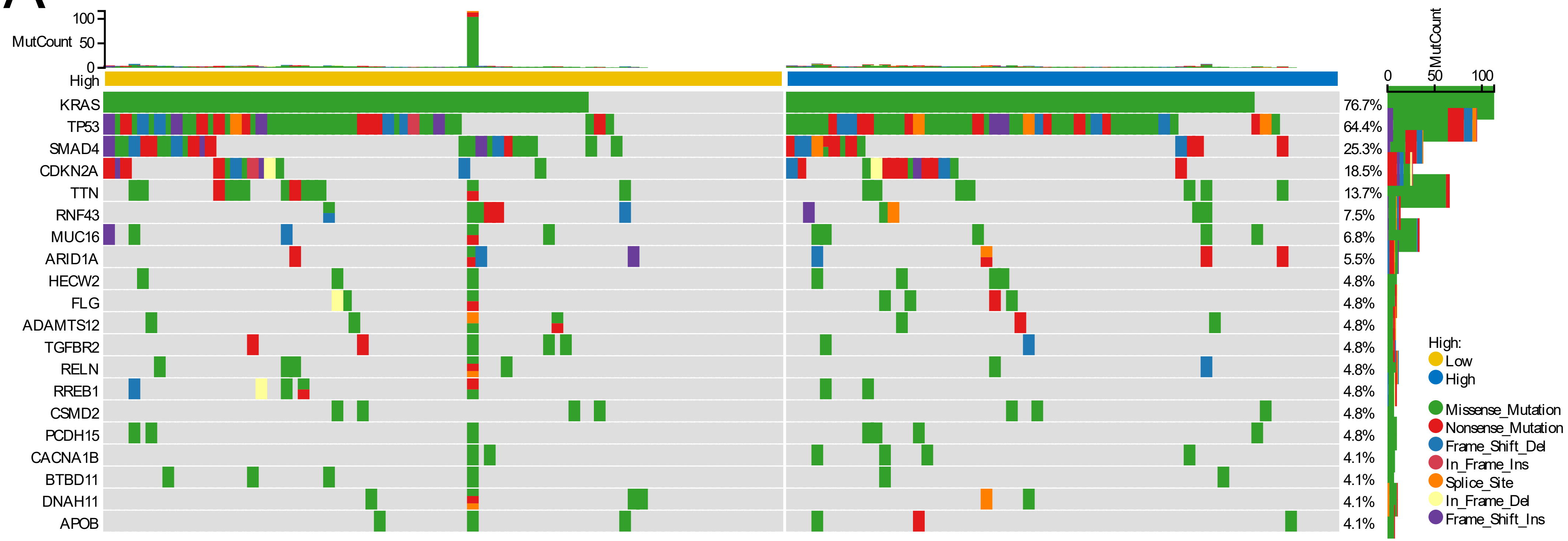
