## Supplementary Tables 1 for "Characterization of Tumor-Associated Endothelial Cells and the Development of a Prognostic Model in Pancreatic Ductal Adenocarcinoma"

Table S1 clinical characteristics of pancreatic ductal adenocarcinoma from multiple cohorts

| Variables | TCGA N=145 | ICGC N=92 | GSE183795 N=139 |
|---|---|---|---|
| Age | | | |
| Median | 65 | 67.5 | - |
| Range | 35-85 | 36-86 | - |
| NA | 0 | 2 | 139 |
| Gender | | | |
| Male | 77 | 47 | - |
| Female | 68 | 44 | - |
| NA | 0 | 1 | 139 |
| TNM Stage | | | |
| I | 12 | 0 | 8 |
| II | 126 | 0 | 105 |
| III | 3 | 0 | 19 |
| IV | 3 | 0 | 6 |
| NA | 1 | 92 | 1 |
| Overall survival | | | |
| Alive | 61 | 32 | 48 |
| Death | 84 | 59 | 86 |
| NA | 0 | 1 | 5 |

*NA, not available.*
