## Supplementary Tables 2 for "Characterization of Tumor-Associated Endothelial Cells and the Development of a Prognostic Model in Pancreatic Ductal Adenocarcinoma"

Table S2 The coefficients of the risk genes in the TaEMS model.

| Gene symbol | coefficient |
| --- | --- |
| DDIT4 | 0.346176247573251 |
| TIE1 | -0.0825796049785623 |
| SEMA6B | -0.378873498317168 |
| PLCB1 | -0.168951734904024 |
| LYZ | -0.301293909944641 |
