## Supplementary Tables 3 for "Characterization of Tumor-Associated Endothelial Cells and the Development of a Prognostic Model in Pancreatic Ductal Adenocarcinoma"

Table S3 Differential sensitivity of 40 drugs group by risk score.

| Drug ID | lower IC50 risk-score group | P-value |
|---|---|---|
| A-770041 | Group Low | 0.028 |
| AKT-inhibitor-VIII | Group Low | 0.0078 |
| AS601245 | Group Low | 0.003 |
| AZ628 | Group Low | 0.00044 |
| AP24534 | Group Low | 0.00047 |
| AZD-0530 | Group Low | 0.00088 |
| AZD8055 | Group Low | 0.00021 |
| Bexarotene | Group Low | 0.033 |
| BMS-509744 | Group Low | 0.0045 |
| BMS-536924 | Group Low | 0.00066 |
| BMS-754807 | Group Low | 0.0018 |
| Bortezomib | Group Low | 0.04 |
| BX-795 | Group Low | 2.9e-06 |
| CEP-701 | Group Low | 0.00063 |
| CHIR-99021 | Group Low | 0.0049 |
| Docetaxel | Group Low | 0.0075 |
| Embelin | Group Low | 0.012 |
| FH535 | Group Low | 0.0027 |
| GDC-0449 | Group Low | 0.0077 |
| GDC0941 | Group Low | 0.028 |
| GSK269962A | Group Low | 0.0021 |
| Imatinib | Group Low | 0.032 |
| KU-55933 | Group Low | 0.0057 |
| MG-132 | Group Low | 0.045 |
| MK-2206 | Group Low | 0.0023 |
| MS-275 | Group Low | 0.02 |
| NU-7441 | Group Low | 0.00084 |
| NVP-BEZ235 | Group Low | 0.0097 |
| NVP-TAE684 | Group Low | 0.00015 |
| PD-173074 | Group Low | 0.00085 |
| PD-0332991 | Group Low | 0.00088 |
| PF-02341066 | Group Low | 3.1e-06 |
| PF-4708671 | Group Low | 0.0059 |
| Pazopanib | Group Low | 0.00076 |
| Roscovitine | Group Low | 0.035 |
| Temsirolimus | Group Low | 0.022 |
| TW-37 | Group Low | 0.0053 |
| Vorinostat | Group Low | 2.5e-05 |
| WH-4-023 | Group Low | 0.006 |
| Sunitinib | Group Low | 0.018 |
